# CORAL: Accurate annotation of compact genomes using long-read RNA-seq, demonstrated in *Oikopleura dioica*

**DOI:** 10.64898/2025.12.04.692336

**Authors:** Nuria P Torres-Aguila, Biel Cassà, Cristian Cañestro

**Affiliations:** Departament de Genètica, Microbiologia i Estadística, Facultat de Biologia, Universitat de Barcelona (UB), Av. Diagonal 645, Barcelona 08028, Spain; Institut de Recerca de la Biodiversitat (IRBio), Universitat de Barcelona (UB), Barcelona, Spain

## Abstract

The burst of high-quality genome assemblies has created an urgent need to improving genome annotation tools, especially for problematic species such as those with compact, fast-evolving genomes in which standard annotation tools underperform when facing short intergenic regions, overlapping UTRs, non-canonical splicing and widespread operons.

We present CORAL (Compact-genome Oriented RNA-based Annotation using Long-reads), a long-read RNA-seq-based workflow for de novo annotation of compact eukaryotic genomes, together with GAMBA (Gene Aggregation tool for Multicistronic Block Annotation), a Rust-based module that detects polycistronic transcriptional units directly from GTF annotations and is integrated within CORAL. Using the exceptionally compact chordate genome of *Oikopleura dioica* as a case study, CORAL increases the number of annotated genes, improves gene-model completeness and reduces chimeric predictions relative to existing annotations, while GAMBA recovers thousands of operons, most of which are independently supported by splice-leader data. Together, CORAL and GAMBA provide an accurate, scalable framework for annotating compact, structurally atypical genomes using long-read transcriptomic data alone.

**Availability and Implementation:** CORAL and GAMBA are open source and available at https://github.com/EvoDevoGenomics-UB/CORAL and https://github.com/nurie05/gamba-tool.

## 1. Introduction

The development of cutting-edge long-read sequencing technologies, such as Oxford Nanopore and Pacific Biosciences, has dramatically increase our capacity to generate high-quality genome assemblies, even at the chromosome scale (van Dijk *et al*., 2023). These advances have enabled the recent emergence of ambitious efforts to sequence all life on Earth, as the Earth BioGenome Project (Lewin *et al*., 2018) and the Darwin Tree of Life (Blaxter *et al*., 2022). However, this burst of newly sequenced genomes has created an urgent need to improving genome annotation tools, especially for problematic species far related to classical model organisms.

A notable example where standard bioinformatic pipelines underperform is in the annotation of compact genomes, especially in fast evolving organisms whose genomic architecture has diverged substantially from canonical patterns (Denoeud *et al*., 2010). Features such as short introns, high gene density with minimal intergenic regions or even intron-nested genes, overlapping UTRs, non-canonical splice sites, and in some cases the unusual organization of genes into polycistronic transcription operon-like units with the often occurrence of transplicing are some of the challenges that need to be addressed for an accurate annotation of compact genomes (Williams *et al*., 2005; Danks *et al*., 2015).

The current approaches for gene annotation include *ab inito* prediction tools, evidence-driven prediction methods, and evidence-based annotation methods (Paniagua *et al*., 2025)*. Ab initio* gene predictions (e.g. AUGUSTUS) rely on statistical models (e.g. Hidden Markov Models) and on stereotypical DNA sequence features (e.g. coding potential of Open Reading Frames, GC content, canonical splice sites, and the presence of start and stop codons), often incorporating algorithm model training with biological data (Stanke *et al*., 2008). Evidence-driven predictions methods (e.g. BRAKER) infer gene models by integrating external biological evidence (e.g. RNA-seq reads, full-length transcripts, ESTs, protein homology) to guide and refine their prediction algorithms (Hoff *et al*., 2019). Evidence-base annotation methods (e.g. StirngTie) rely exclusively on biological data to reconstruct gene models with high accuracy, although their performance is limited by the availability and quality of such evidence (Kovaka *et al*., 2019).

In most cases, the biological evidence used for gene annotation consists on short-read RNA-seq data, which is characterized by high coverage and low error rates but it is limited in its ability to resolve transcript isoforms. Moreover, incomplete or unreliable biological evidence can hinder the correct identification of non-canonical splice sites and, consequently, affect overall gene structure accuracy.

Long-read RNA-seq represents a powerful alternative, as each long-read molecule captures with high precision the gene structure and directly reveals transcript variants. Annotation based on long-reads, however, can be challenging in organisms containing operons, where full-length unprocessed operon transcripts may be mistakenly interpreted by annotation tools as a single gene with multiple isoforms rather than distinct genes arranged in an operon.

To address the challenges for an accurate annotation of compact genomes we have developed CORAL (Compact-genome Oriented RNA-based Annotation using Long-reads), a long-read transcriptome-driven annotation pipeline designed for compact eukaryotic genomes. This bioinformatic workflow uses long-read RNA-seq data to annotate compact genomes without using any prediction tool and circumventing issues associated with short intergenic regions, overlapping UTRs, operons and other features that challenge conventional annotation approaches. To specifically address operon annotation, we have developed GAMBA (Gene Aggregation tool for Multicistronic Block Annotation), a Rust-based tool that identifies polycistronic transcriptional units directly from GTF annotations and integrated within CORAL. To demonstrate the function and accuracy of CORAL we used *Oikopleura dioica* as case study.

*O. dioica* is a fast-evolving chordate with an extremely compact genome (Seo *et al*., 2001; Bliznina *et al*., 2021). Its 70 Mb genome, smaller *than Caenorhabditis elegans* or *Drosophila* genomes, contains around 16,000 annotated genes, resulting in a very high gene density with short intergenic regions ranging from 50 bp up to few kilobases (Seo *et al*., 2001; Bliznina *et al*., 2021). This high density is accompanied by overlapping UTRs, frequent non-canonical splice sites, and abundant operons (Ganot *et al*., 2004; Denoeud *et al*., 2010; Danks *et al*., 2015; Ferrández-Roldán *et al*., 2019; Frey and Pucker, 2020). Genomic complexity is further increased by extensive gene loss, lineage-specific duplications, and chromosomal rearrangements such as inversions and translocations (Martí-Solans *et al*., 2021; Masunaga *et al*., 2022; Plessy *et al*., 2024; Sánchez-Serna *et al*., 2025). Comparative genomic studies among *O. dioica* populations worldwide revealed a massive genome scrambling, suggesting the presence of cryptic species (Masunaga *et al*., 2022; Plessy *et al*., 2024). Together, these features introduce multiple layers of variability that make the annotation of *O. dioica*’s genome particularly challenging.

Here, we demonstrate how CORAL overcomes the challenges of the compact and fast-evolving genome of *O. dioica*, delivering an accurate and biologically supported annotation. CORAL produces a higher number of annotated genes, fewer chimeric gene models, richer sets of alternative-splice variants, and robustly define operons, all directly supported by long-read RNA evidence. These results position CORAL as a robust framework for annotating compact, fast-evolving, or structurally atypical eukaryotic genomes.

## 2. Material and Methods

### 2.1. Overview of the CORAL workflow

CORAL (Compact-genome Oriented RNA-based Annotation using Long-reads; Figure 1) is a custom workflow developed with Snakemake (Köster and Rahmann, 2012; Mölder *et al*., 2021) for the annotation of eukaryotic compact genomes using long-read RNA-seq data. The user specifies the datasets to be included and assigns them a sample ID to each. If needed, multiple datasets can share the same sample ID.

**Figure 1.**
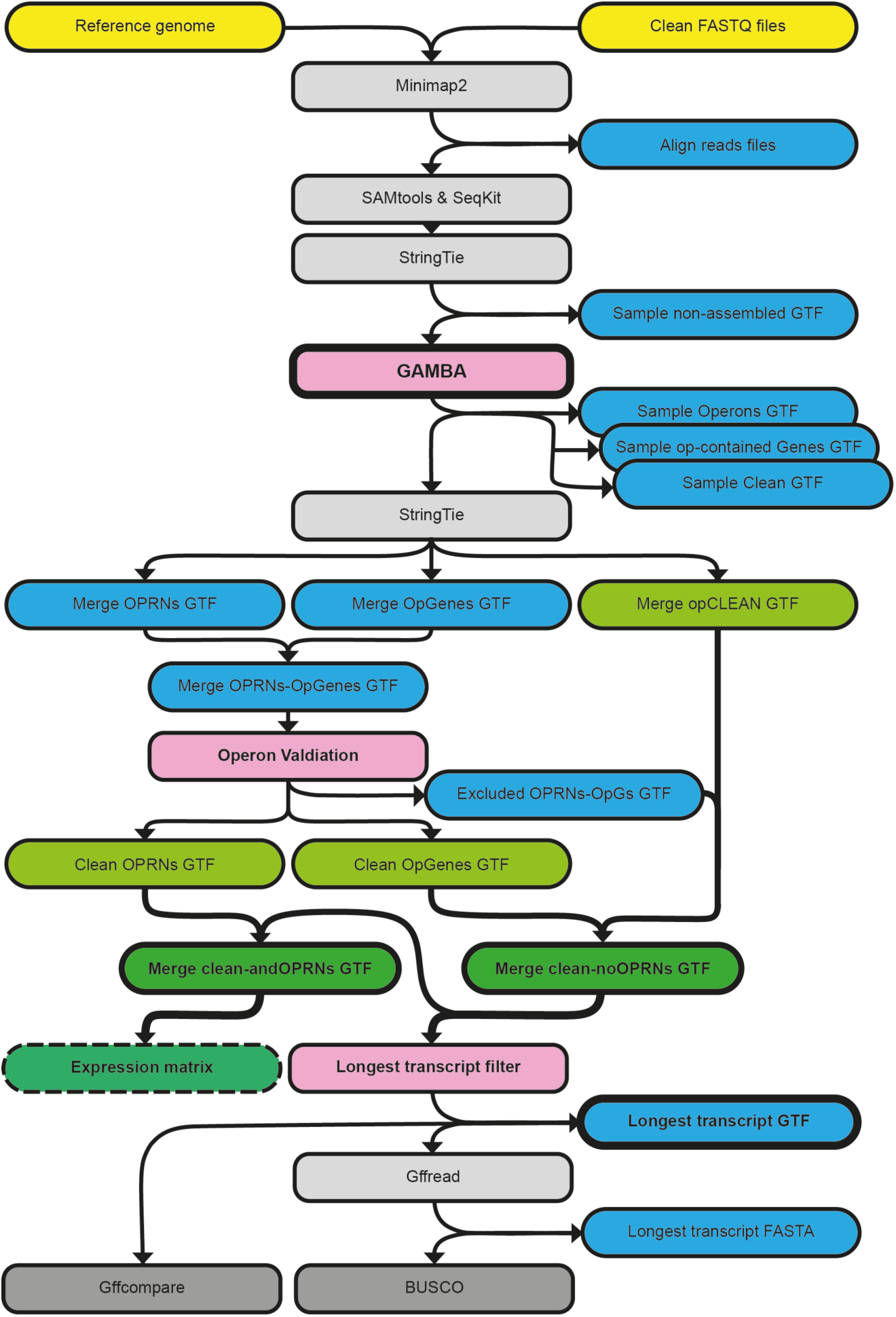
**Schematic of the CORAL workflow**. This annotation workflow for compact genomes uses only biological data (i.e. long-read RNA-seq data) to annotate genes and operons. Yellow: input files; Light grey: bioinformatic tools; Pink: novel bioinformatic tools developed; Dark grey: bioinformatic tools used for quality check; Blue: intermediate output files: Light green: preliminary annotation files; Dark green: final annotation files on GTF format; Dashed box: optional expression matrix output.

The workflow process starts with the alignment of QC-passed long-read sequences to the provided reference genome with minimap2 (Li, 2021, 2018). On this step, the user is allowed to set parameters for the index creation and for the alignment process on the configuration file. The ‘-G’ parameter, that set the maximum intron size allowed during the mapping, is explicitly used and it is required to be set by the user on the configuration file.

After alignment, a formatting and filtering process is performed with SAMtools (Danecek *et al*., 2021), and SeqKit (Shen *et al*., 2016), respectively. Filtering is performed with the ‘-q’ option of the SeqKit ‘bam’ module, applying a user-defined minimum mapping quality threshold. Subsequently, preliminary annotations are generated for each sample ID using StringTie (Kovaka *et al*., 2019) (Figure 1: ‘Sample non-assembled GTF’). The ‘-L’ and ‘-R’ parameters are applied to enable long-read-specific processing and to prevent chimeric transcript assembly. Strand orientation is specified in the CORAL configuration file, which also allows users to supply any other StringTie parameters as needed.

Then, for each sample ID annotation, CORAL executes our developed tool GAMBA (Gene Aggregation tool for Multicistronic Block Annotation). This Rust-based tool identifies polycistronic transcriptional units (operons) directly from GTF annotation files. Specifically, GAMBA clusters adjacent transcripts based on their genomic position, structure, and expression levels (i.e. coverage and FPKM), enabling the reliable delineation of multicistronic blocks supported by biological evidence.

A transcript is classified as putative operon transcript when it contains other transcripts (with a tolerance of 250 bp and on the same strand) and satisfies the following criteria: (i) it contains at least two non-overlapping transcripts; (ii) the contained transcripts have a higher coverage than the container transcript multiplied by a user-defined threshold (default: 1); (iii) the contained transcript overlaps at exonic level with the container transcript; and (iv) the container transcript is not contained within a higher coverage transcript. These criteria ensure that detected operons are supported structurally and by expression evidence. When two contained transcript overlaps (defined as overlap of ≥50 bp), GAMBA retains the transcript with the highest FPKM to avoid redundancy on contained transcripts, prioritizing multi-exonic transcripts over mono-exonic ones when applicable. GAMBA generates three output annotation files (Figure 1): (1) ‘Operons GTF’, containing the transcripts identified as operons; (2) ‘op-contained Genes’, containing the transcripts identified as operon-contained gene; and (3) ‘Clean GTF’, containing all non-operon-related transcripts after filtering out poor-evidence models (coverage ≤1 for multi-exonic transcripts, and ≤10 for mono-exonic transcripts).

After running GAMBA, CORAL generates consensus annotations for three categories independently: operons, operon-contained genes and non-operon genes. This step uses the ‘--merge’ funtion of StringTie (Figure 1: ‘Merge OPRNs GTF’, ‘Merge OpGenes GTF’, and ‘Merge opCLEAN GTF’). For operons and operon-contained genes, CORAL uses the parameters ‘-F’, ‘-T’ and ‘-c’ set to 0 to prevent the exclusion of any transcript supported by the long-read data. For non-operon genes these parameters are user-defined on the configuration file. The ‘-l’ parameter is automatically set to ‘OPRNs’, ‘OpG’, or ‘g’ to label the merged annotations for operons, operon-contained genes, and non-operon genes, respectively. The ‘-g’ parameter is user-defined, independently, for operon-contained genes and non-operon genes, but is fixed to 0 during operon annotation to favour the assembly of full-length operonic transcripts.

Next, a guide annotation is created by merging the operon annotation and the operon-contained-gene annotation (Figure 1: ‘Merge_OPRNs-OpGs’). This is done by concatenating both GTF files with the ‘cat’ command, followed by sorting with gffread (Pertea and Pertea, 2020) using the ‘--sort-alpha’ option. This resulting guide annotation is then validated with a custom Python script (operon_validation.py) which removes any false positive operon (i.e. operon containing a single gene after the consensus merge) along with its contained genes (Figure 1: ‘Excluded OPRNs-OpGs GTF’). The script outputs a summary file reporting the number of OPRNS and OpGs identified, along with clean consensus annotations for both operons and operon-contained genes (Figure 1: ‘Clean OPRNs GTF’ and ‘Clean OpGenes GTF’).

The final merged annotation is generated using the StringTie ‘--merge’ function, combining the consensus annotation of non-operon genes with the discarded false positive operons and operon-contained genes (Figure 1: ‘Merge clean-noOPRNs GTF’). In this step, CORAL uses the ‘-G’ parameter with the ‘Clean OpGenes GTF’ file as guide annotation, sets the ‘-F’, ‘-T’ and ‘-c’ parameters to 0 to retain all transcript models, assigns ‘-l g’ for labelling proposes, and allows the used to define the ‘-f’ and ‘-g’ parameters. Finally, a comprehensive annotation that includes polycistronic transcripts is created by appending the consensus operon annotation of the final consensus annotations (Figure 1: ‘Merge clean-andOPRNs GTF’). This is performed using the ‘cat’ command followed by gffread with the ‘--sort-alpha’ option.

The final merged annotation is processed with a custom Python script (Longest_transcript_filter.py) to obtain a non-redundant annotation which contains only a single transcript per gene selecting the longest one (Figure 1: ‘Longest transcript GTF’). This non-redundant annotation is used to assay the quality of the obtained annotation using BUSCO (Manni *et al*., 2021) and, if a reference annotation is provided, gffcompare (Pertea and Pertea, 2020).

As optional feature, CORAL can create an expression matrix that includes, for each sample ID specified, the raw reads seen for each transcript or gene of the final obtained annotation, a valuable information to study expression dynamics. This matrix is done following the protocol specified for use StringTie with DESeq2 (Pertea *et al*., 2016).

### 2.2. Biological material and RNA extraction

*O. dioica* specimens were acquired from the animal cultures maintained in the laboratory (Martí-Solans *et al*., 2015). A total of twelve different stages of *O. dioica* lifecycle were selected to provide representative coverage of each phase of development. The selected stages were: egg, 8-cells, 32-cells, 64-cells, incipient tailbud (ITB), mid tailbud (MTB), early hatchling (EH), mid hatchling (MH), late hatchling (LH), adult-juvenile, pre-mature female, and male. For collecting pre-adult developmental stages *in vitro* fertilizations were performed following published protocol (Martí-Solans *et al*., 2015). RNA extractions were done using QIAGEN kit RNeasy Plus Micro kit (ref. 74104). Samples, containing at least 100 embryos or 20 adult animals, were added directly to the lysis buffer of the kit to start the extraction protocol. Obtained RNA was stored at-80 °C util use for sequencing.

### 2.3. Long-read data processing and mapping optimization

Long-read RNA-seq data was obtained using Oxford Nanopore Technologies (ONT) protocol PCR-cDNA with BARCODING in a GridION sequencer with the base-calling algorithm ‘Dorado’ (version 7.4.12+0e5e75c49). All samples were processed in a single run, except for sample BP10 that was sequenced independently. Quality Controls (QC) of the data included Pychopper (ONT) and SeqKit (Shen *et al*., 2016), to trim the adapters and remove short sequences (under 200bp), respectively.

Different-G parameters ranging from 2kb to 50kb were tested to select the best parameter for mapping. The one showing more aligned reads was selected. This test was done in a single sample: BP10, corresponding to the stage ‘adult-juvenile’, which contained more reads.

### 2.4. *O. dioica* genome annotation with CORAL

A *de novo* annotation of *O. dioica* genome was done using the 12 QC-processed long-read datasets as input for CORAL. The reference genomes used were Bar2p4 (Plessy *et al*., 2024) for the Barcelona (Spain) population, and ASM20953v1 (Denoeud *et al*., 2010) for the Bergen (Norway) population.

For alignment, minimap2 (v2.30) was executed with the parameters specified in the CORAL configuration file: ‘-k14’, for indexing, and ‘-ax splice’ with a maximum intron length of ‘10k’ for the alignment step. Preliminary sample annotations were produced with StringTie (v 3.0.3), using the configuration-file settings: ‘--fr’, for strand orientation, and ‘-M 0.75-j 2-a 15’ as additional parameters, to improve junction accuracy and limit multimapping. Operons (OPRNs) and operon-contained genes (OpGs) were identified by GAMBA (v2.0). For the OpGs consensus annotation the ‘-g’ parameter was set to ‘-150’ to prevent gene fusion. For the non-operon genes consensus annotation and the final merged annotation, the ‘-f’ parameter was set to 0.2 to reduce transcript redundancy, the ‘-g’ parameter was set to ‘-60’ to reduce computational load, and the optional parameters ‘-F 0.5-T 0 - c 2’ were provided to increase transcript robustness. All consensus and merged annotations were generated with StringTie (v3.0.3), with all parameters specified in the CORAL configuration file. Annotation quality was assessed with BUSCO (v5.8.2) and gffcompare (v0.12.10). The reference annotations used for gffcompare correspond to those reported in Plessy et al., 2024.

### 2.5. Validation of operons with CAGE data

To further validate the operons detected by GAMBA, we used the splice-leaders (SL) defined previously on the Bergen *O. dioica* genome (Danks et al., 2015).To confirm that detected OPRNs and OpGs on the Bergen transcriptome had SL as expected we formatted the SL data into a GTF file and used a custom python script (‘sl_validation.py’) to check for SL sites on the 5’ of the described OpGs.

## 3. Results

### 3.1. Long-read data and mapping optimization

We collected and sequenced, using Oxford Nanopore Technologies (ONT), the total RNA of 12 different stages of *O. dioica*: eggs, 8-cells, 32-cells, 64-cells, incipient tailbud (ITB), mid tailbud (MTB), early hatchling (EH), mid hatchling (MH), late hatchling (LH), adult-juveniles, pre-mature females, and males. On average, a total of 10.7M reads were obtain per samples, from which an 83.9% pass our quality controls (QCs) having an average of 9M reads per sample (Table 1).

**Table 1.** Sequencing statistics per sample. QC: quality control; Min.: minimum; Avg.: average; Max.: maximum; Len.: length; iTB: incipient tailbud: midTB: mid tailbud; JH: just hatchling; MH: mid hatchling; LH: late hatchling; Pre-females: premature females; M: million reads; Gb: giga-bases.

| Sample | Stage | Raw reads | %QC pass | Clean reads | Min. Len. | Avg. Len. | Max. Len. | Sum Len. |
| --- | --- | --- | --- | --- | --- | --- | --- | --- |
| BP01 | Egg | 10.77M | 84.4% | 9.08M | 200 | 862.4 | 8353 | 7.83 Gb |
| BP02 | 8-cells | 7.59M | 85.9% | 6.52M | 200 | 888.9 | 8209 | 5.80 Gb |
| BP03 | 32-cells | 10.96M | 83.7% | 9.18M | 200 | 847.9 | 6014 | 7.78 Gb |
| BP04 | 64-cells | 9.22M | 84.9% | 7.83M | 200 | 861.7 | 5824 | 6.74 Gb |
| BP05 | iTB | 6.74M | 84.1% | 5.66M | 200 | 870.3 | 8372 | 4.93 Gb |
| BP06 | midTB | 7.69M | 84.3% | 6.48M | 200 | 840.7 | 8378 | 5.45 Gb |
| BP07 | JH | 8.23M | 84.3% | 6.94M | 200 | 855.1 | 8363 | 5.93 Gb |
| BP08 | MH | 10.35M | 84.5% | 8.75M | 200 | 830.7 | 9159 | 7.26 Gb |
| BP09 | LH | 10.04M | 84.2% | 8.45M | 200 | 825.6 | 7404 | 6.98 Gb |
| BP10 | Juveniles | 24.79M | 81.3% | 20.17M | 200 | 787.2 | 9194 | 15.87 Gb |
| BP11 | Pre-females | 12.12M | 83.1% | 10.07M | 200 | 926.5 | 7324 | 9.33 Gb |
| BP12 | Males | 10.72M | 82.2% | 8.81M | 200 | 1003.3 | 5948 | 8.84 Gb |
| **AVG.** | **---** | **10.77M** | **83.9%** | **9.00M** | **200** | **866.7** | **7711.8** | **7.73 Gb** |

Among the different samples, BP10 (adult-juveniles) was selected for mapping optimization, as it showed the higher number of reads after QCs. To optimize the mapping on compact genomes, we tested different values of the minimap2 parameter ‘-G’, which sets the maximum intronic size allowed during the mapping process. Considering that the described introns of *O. dioica* raged from 50 bp up to few kilobases (Seo *et al*., 2001; Bliznina *et al*., 2021), we tested: 2K, 5K, 10K, 20K, and 50K.

Results showed that, despite having a high number of reads mapped in all cases, the total number of reads mapped was slightly different (Table 2). Setting the parameter ‘-G’ to 10k showed the higher percentage of mapped reads: 98.73%, 7,409 more reads than the second best one. Therefore, the ‘-G’ parameter was set to 10k to map all the other samples (Table 3), mapping on average 98.6±0.2% of the reads.

**Table 2.** Variation of alignment results when modifying ‘-G’ parameter on *minimap2*. Alignment results of sample BP10 (Juveniles) for diUerent maximum intronic sizes allowed during the mapping into the Barcelona genome, ranging from 2Kb up to 50Kb.

| ‘-G’ parameter | #Aligned reads | %Aligned reads |
| --- | --- | --- |
| 2k | 19,900,881 | 98.65% |
| 5k | 19,904,816 | 98.67% |
| 10k | 19,916,874 | 98.73% |
| 20k | 19,908,222 | 98.69% |
| 50k | 19,909,465 | 98.69% |

**Table 3.** Alignment statistics of each sample. Alignment statistics after mapping with *minimap2* (using ‘-G 10k’) on the Barcelona and Noth Sea genomes. AVG.: average. M: million reads.

| Sample | Clean reads | Barcelona<br>Aligned reads | Bergen<br>Aligned reads |
| --- | --- | --- | --- |
| BP01 | 9.08M | 8.97M (98.7%) | 8.70M (95.8%) |
| BP02 | 6.52M | 6.45M (99.0%) | 6.26M (95.9%) |
| BP03 | 9.18M | 9.10M (99.1%) | 8.83M (96.2%) |
| BP04 | 7.83M | 7.75M (99.0%) | 7.53M (96.2%) |
| BP05 | 5.66M | 5.59M (98.6%) | 5.41M (95.4%) |
| BP06 | 6.48M | 6.38M (98.5%) | 6.14M (94.7%) |
| BP07 | 6.94M | 6.86M (98.8%) | 6.56M (94.5%) |
| BP08 | 8.75M | 8.64M (98.8%) | 8.40M (96.0%) |
| BP09 | 8.45M | 8.34M (98.7%) | 8.10M (95.8%) |
| BP10 | 20.17M | 19.91M (98.7%) | 19.20M (95.2%) |
| BP11 | 10.07M | 9.98M (99.1%) | 9.67M (96.0%) |
| BP12 | 8.81M | 8.51M (96.5%) | 8.47M (96.0%) |
| AVG. | **9.00M** | **8.87M (98.6%)** | **8.61M (95.6%)** |

### 3.2. Operon identification

Using our custom tool GAMBA, we were able to identify polycistronic transcripts in all the samples used. On average 814±142 operons (OPRNs), with 1,877±348 operon-contained genes (OpGs), were identified per sample on the Barcelona genome, most of them composed of two genes (Figure 2A). After merging the identified operon-related transcripts into a consensus annotation, the resulting annotation contained 2,434 OPRNs with a total of 6,885 OpGs. Of the identified OPRNs, 61% were composed for 2-genes, 19% for 3-genes, 9% of 4-genes, 5% of 5-genes and 6% of over 5-genes (Figure 2A). The OPRNs with most genes identified were OPRN.1403 and OPRN.1684 with 12 genes, and OPRN.1760 with 13 genes, all located on the long arm of chromosome PAR (loci PAR:7,654,298-7,671,778, PAR:11,697,313-11,714,400, and PAR:13,009,381-13,023,233, respectively).

**Figure 2.**
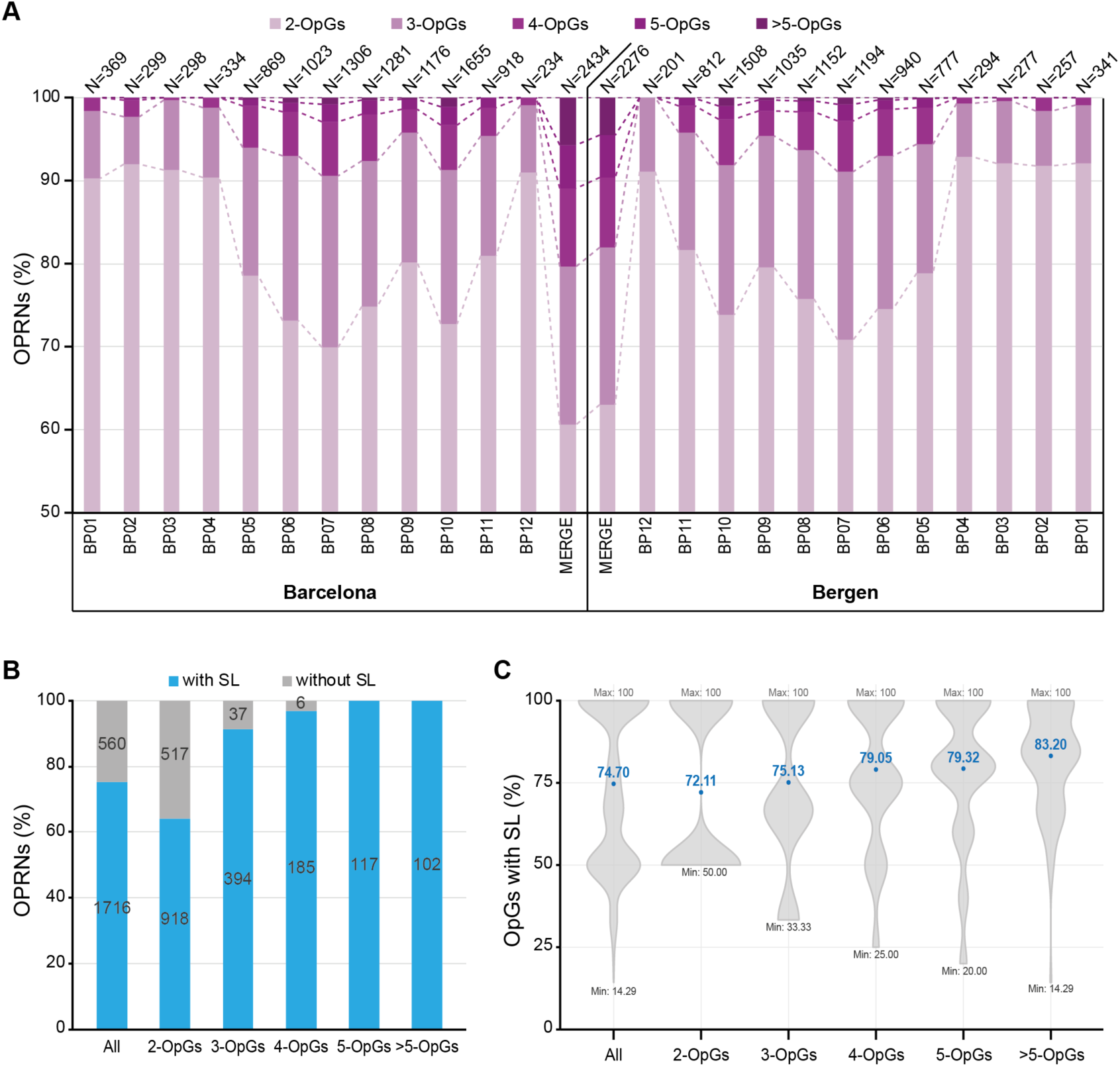
Operon metrics across *O. dioica* genomes. A) Percentage of operons (OPRNs) found according to number of genes contained (OpGs), on each sample (BP01-BP12) and on the merged annotation (Merge), for the Barcelona and the Bergen *O. dioica* genomes. Notice that Y-axis start on 50%. N: total number of operons. B) Percentage of operons (OPRNs), found on the Bergen *O. dioica* genome, that contained at least one splice-leader site (SL). Numbers are shown for the whole set of OPRNs (All) and divided by the number of genes contained (OpGs categories). C) Percentage of genes within an OPRN with SL sites, that had a SL site at 5’end for the whole set of OPRNs with SL sites (All) and divided by the number of genes contained (OpGs categories). Mean (dot) and its value are shown in blue for each category. Min: minimum value observed; Max: maximum value observed.

Similar results were found when using the Bergen reference genome (Figure 2A), with on average 732±129 OPRNs and 1680±314 OpGs per sample and a total of 2,276 OPRNs with 6,226 OpGs. Furthermore, the Bergen operon containing more genes was the same as the identified on the Barcelona genome, OPRN.1760, named OPRN.1711 on the Bergen annotation with 13 genes and 14kb long (locus scaffold_49:25,751-39,181).

Interestingly, a previously described operon of 9 genes (locus PAR:8,766,290-8,785,222 on the Barcelona genome) conserved within *O. dioica* populations, was not found as operon in our data. There was no evidence of long-reads supporting the operon fully. In contrast, we only found at that locus a 3-gene composed operon, found at the beginning, and a 2-gene composed operon, on the middle.

This highlights the need to validate predicted operons with biological data.

To validate the accuracy of our GAMBA annotation tool, which is based solo on long-read data, we aimed to verify the OPRNs and OpGs found using available SL data (Danks *et al*., 2015). Our results showed that 75.4% (1,716) of the found Bergen OPRNs contained SL sites on 5’ end positions of their contained OpGs (Figure 2B). Within those, on average 74.7% of their contained OpGs had a SL site at the 5’ end (Figure 2C), this percentages increased sequentially when divided the OPRNs by number of contained OpGs (Figure 2B-C).

### 3.3. Annotation improvement

The consensus annotation created for the Barcelona genome contained 18,826 genes, which is a 23% increase (3,486 more genes) on the total number of genes available on the reference annotation, with 22,691 transcripts without the operon representative transcripts, and 26,706 when including the operon transcripts (Figure 3A). Surprisingly, 9% of transcripts corresponded to wrongly predicted genes on the reference annotation (categories ‘o’ and ‘x’; Figure 3B), and an additional 13% were completely novel transcripts (categories ‘i’ and ‘u’; Figure 3B). Only 39% of the annotated transcripts were perfectly matching the reference annotation (categories ‘=’, ‘c’ and ‘k’; Figure 3B), with an additional 38% corresponded to potential isoform variants (categories ‘j’, ‘m’ and ‘n’; Figure 3B).

**Figure 3.**
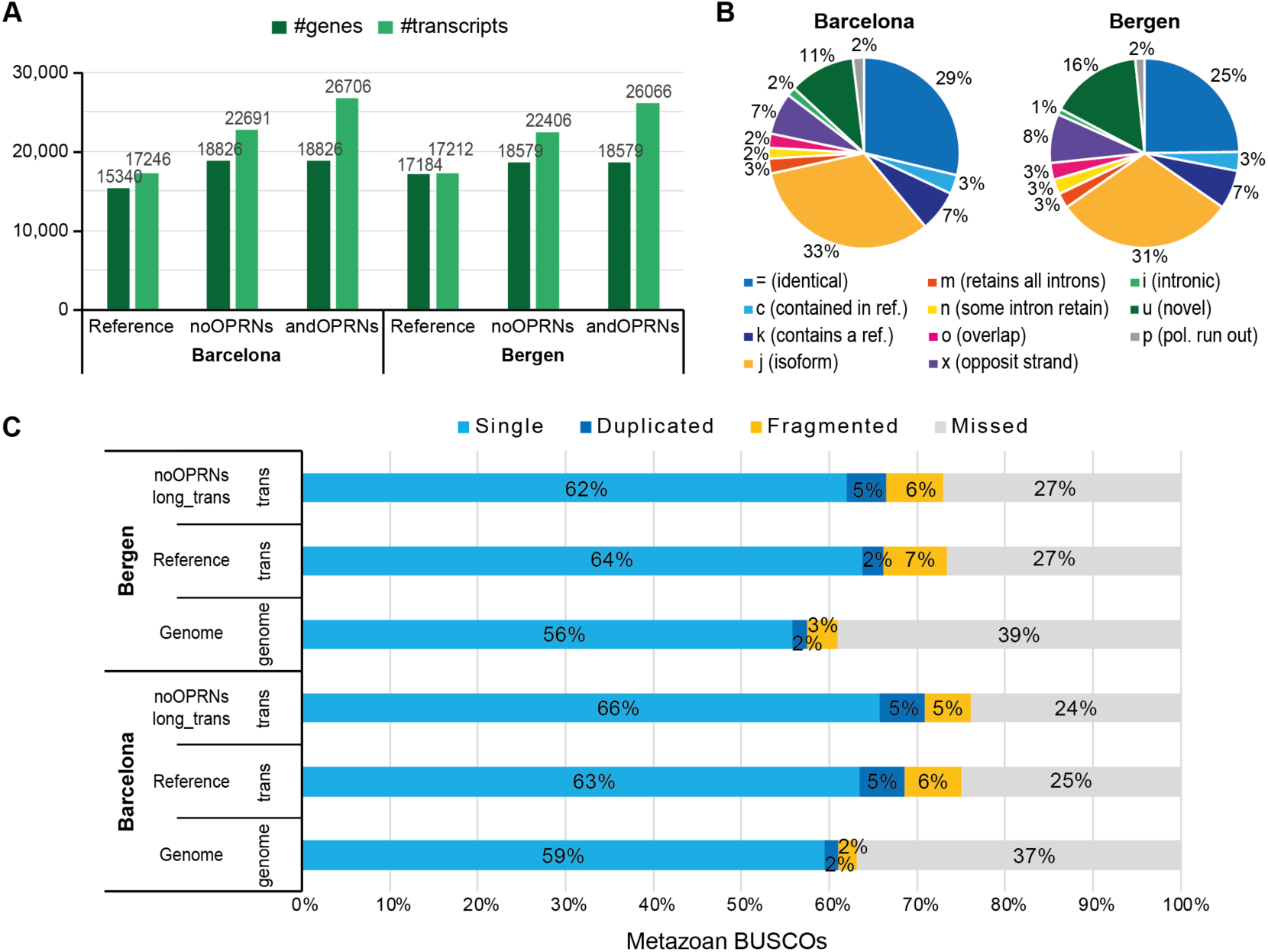
Annotation metrics across *O. dioica* genomes. A) Gene and transcript content of the reference annotation (Reference) and the novel annotation (noOPRNs: includes gene and operon-contained gene transcripts, andOPRNs: includes gene, operon-contained gene and operon transcripts). B) Pie chart of the transcript type classification by *g.compare* when comparing the novel annotation and the reference annotation. = (blue), identical reference-query match; c (light-blue), complete query match within reference; k (dark-blue), complete reference match within query; j (gold), some splice site mismatch (potential novel isoform); m (orange), all intron retained; n (yellow), some introns retained; o (magenta), partial overlapping match; x (purple), exon match on opposite strand; i (light-green), contained within reference intron; u (gray), no match of query within reference (novel). Categories e (single exon match), y (reference contained within intron), s (reference contained within intron), and r (repeat) were excluded as represented less than 1% of the transcripts. C) Percentage of metazoan BUSCOs found on the annotations and on the genome, using the corresponding mode ‘trans’ or ‘genome’. Blue: Single complete BUSCOs; Dark-blue: duplicated complete BUSCOs; Yellow: Fragmented BUSCOs; Gray: Missing BUSCOs.

Furthermore, the BUSCO analysis showed an increase of 2.3% in the number of Complete BUSCOs, with a reduction on Fragmented and Missing BUSCOs (1.3% and 1.0% reduction respectively; Figure 3C). Similar results were found when using the Bergen genome (Figure 3A-C), despite the increase on the total number of genes annotated (8,1% more genes) and the increase of Complete BUSCOs (0.3% increase) were more discreet.

## 4. Discussion

*O. dioica* is characterized for the most compact genome known among chordates (Seo et al., 2001). Its extreme compactness, together with extensive gene losses, lineage-specific duplications, and structural rearrangements, has made accurate annotation challenging, with published gene sets presenting low BUSCO completeness (<70%; (Plessy *et al*., 2024). Here we presented a substantial improvement over the current reference annotations, generated using a previously trained AGUSTUS model. CORAL, by using only long-read RNA-seq data, increased the total number of annotated genes on both populations examined (Barcelona and Bergen), despite one presenting a highly fragmented genome assembly. The resulting *de novo* annotations achieved 71% of complete BUSCOs on the Barcelona genome and 67% on the Bergen genome, highlighting the robustness and assembly-independence of the CORAL workflow.

A key advantage of CORAL is the use of strand-specific long-read transcriptomic data, which enabled the detection of genes transcribed from opposite strands within the same genomic regions. Such opposite-strand or antisense gene arrangements are commonly underrepresented in *ab initio* pipelines (e.g., AUGUSTUS, BRAKER), which usually retain only the highest-scoring model per locus. By relying exclusively on empirical transcriptional evidence, CORAL captured both sense and antisense gene models, improving gene-structure accuracy and better reflecting the biological complexity of compact genomes.

By using GAMBA, we were able to identify 2,434 and 2,276 polycistronic transcriptional units in the Barcelona and Bergen genomes, containing 6,786 and 6,226 genes, respectively. In addition to the intrinsic transcriptional evidence of CORAL, in the Bergen genome, 75.4% of the found operons were additionally supported by SL-mapping data, with ∼74.7% of OpGs displaying a SL site at 5’end. This is consistent with the expectation of the first gene on an operon typically lacks a SL sequence. These results represent the first genome-wide validation of operon organization in *O. dioica* relying exclusively on transcriptomic data, positioning GAMBA as a valuable tool for investigating operon architecture, expression dynamics and regulatory mechanisms in eukaryotes.

In conclusion, our results demonstrate that CORAL is a robust and assembly-independent workflow for annotating compact eukaryotic genomes using exclusively biological data. By exploiting the advantage of the long-read sequencing, CORAL accurately reconstructs gene models, resolves antisense transcription, and robustly identifies operonic architectures. Together, CORAL and GAMBA provide a robust, scalable, and biologically informed strategy for annotate challenging eukaryotic genomes using solely long-read RNA-seq data.

## Acknowledgments

The authors thank the ‘Centres Científics i Tecnològics de la UB’ for sea water supply, Novogene for their sequencing services and, especially, Sebastian Artime Paoletti for running the *Oikopleura* facility at the University of Barcelona, Nicholas M. Luscombe’s lab members for helpful discussions, Josep F. Abril Ferrando for his support on our computational server at the UB, and Miguel Vera Belmonte for his assistance with Rust code.

## Author contributions

Conceptualization, C.C. and NP.T.; Methodology, NP.T.; Software, NP.T.; Validation, NP.T. and B.C.; Formal Analysis, NP.T.; Investigation, NP.T.; Resources, NP.T. and B.C.; Data Curation, NP.T.; Writing – Original Draft Preparation, NP.T.; Writing – Review & Editing, C.C. and NP.T.; Visualization, NP.T.; Supervision, C.C.; Project Administration, NP.T. and C.C.; Funding Acquisition, NP.T. and C.C.

## Data Availability

*O. dioica* long-read RNA-seq data generated is available at the Sequence Read Archive (SRA) database under accession numbers SRR35841405-SRR35841416, BioProject PRJNA1347569. The annotation files produced are deposited on Zenodo (DOI: <u>10.5281/zenodo.17791010</u>). The SL CAGE data used, from Danks et al., 2015, was downloaded from the GEO database, accession number GSE50849. All the scripts used for data processing are available in GitHub, repository: https://github.com/EvoDevoGenomics-UB/CORAL and https://github.com/nurie05/gamba-tool.

